# Integrative Genomic Analyses Reveal Putative Cell Type-specific Targets of the *Drosophila* Ets Transcription Factor Pointed

**DOI:** 10.1101/2023.09.08.556887

**Authors:** Komal Kumar Bollepogu Raja, Kelvin Yeung, Yoon-Kyung Shim, Graeme Mardon

## Abstract

The Ets domain transcription factors direct diverse biological processes throughout all metazoans and are implicated in development as well as in tumor initiation, progression and metastasis. The *Drosophila* Ets transcription factor Pointed (Pnt) is required for several aspects of eye development and regulates cell cycle progression, specification, and differentiation. Despite its critical role in development, very few targets of Pnt have been reported previously. Here, we used chromatin immunoprecipitation with high-throughput sequencing (ChIP-seq) to determine the genome-wide occupancy of Pnt in late larval eye discs. We identified enriched regions that mapped to an average of 6,941 genes, the vast majority of which are novel putative Pnt targets. Integrating ChIP-seq data with two other larval eye single cell genomics datasets (scRNA-seq and snATAC-seq) reveals genes that may be putative cell type-specific genes regulated by Pnt. Finally, our ChIP-seq data predict cell type-specific functional enhancers that were not reported previously. Our study provides a greatly expanded list of putative Pnt targets in the eye and is a resource for future studies that will allow mechanistic insights into complex developmental processes regulated by Pnt.

## Introduction

Precise orchestration of complex biological processes such as cell signaling and gene expression is necessary for normal development of organisms. Gene expression is an integral part of all organisms and transcription factors (TFs) play vital roles in this process. TFs are DNA-binding proteins that regulate the activity of genes and are key determinants of gene function at a given time in development. Further, TFs are involved in a wide range of cellular processes as well as in mechanisms underlying response to stress, disease and apoptosis. Therefore, analyzing the activity of TFs and the target genes they regulate will help unravel their significance in development and disease.

The E26 transformation specific (Ets) family of TFs are highly conserved in metazoans and ensure proper development by regulating diverse processes such as proliferation, differentiation and apoptosis(1-3). The Ets factors contain a winged helix-turn-helix DNA-binding domain that recognizes a core GGAA/T consensus sequence and serve as transcriptional activators or repressors depending on the context. Furthermore, additional transcription factors and the DNA sequences adjacent to the Ets binding sites determine the DNA-binding specificity of Ets factors(4). This combinatorial interaction between Ets factors and other proteins dictate context-dependent regulation and downstream target specificity. Ets TFs function downstream of several signaling pathways such as the Mitogen-activated protein kinase (MAPK) pathway, and their aberrant expression contributes to tumor initiation, progression, and malignant transformation(5-9). Therefore, continued investigation of gene regulation by Ets factors will provide mechanistic insights into why deregulation of these factors results in tumor initiation and growth.

The fruit fly *Drosophila melanogaster* is one of the preferred models to study Ets TFs. Eight Ets family proteins have been identified in *Drosophila*, including Pointed (Pnt)(10,11). Pnt is required during embryonic development and plays a critical role in the specification and differentiation of cells in several tissues, including the ventral ectoderm and the nervous and tracheal systems(10,12-15). In the fly eye, *pnt* regulates the progression of the morphogenetic furrow (MF), cell cycle, and cell differentiation and is required for cell survival(16-21). Each ommatidium in the *Drosophila* eye is a repeating unit of a cluster of 20 neuronal and non-neuronal cells. Differentiation begins at the posterior margin of the second instar larval eye disc with the initiation of the MF that moves anteriorly leaving differentiated cells and mature progenitors behind it. Anterior to the MF, cells are in an undifferentiated state and poised to undergo differentiation. The photoreceptor R8 differentiates first followed by R2/5 and R3/4. Then the remaining undifferentiated cells undergo another round of division known as the ‘second mitotic wave’ (SMW). R1/6, R7 and cone cells differentiate from the cells exiting the SMW. The remaining non-neuronal pigment cells differentiate during pupal stages. Interestingly, with the exception of R8, the reiterative use of the *Epidermal growth factor receptor* (*Egfr*) is required for the sequential differentiation of all cells in the eye(17). In response to *Egfr* signaling, Pnt is phosphorylated, which allows it to function as a transcriptional activator(16). How Pnt functions downstream of Egfr to induce different cell fates in the eye in entirely not clear. However, it has been suggested that Pnt interacts with other factors to regulate the expression of different target genes depending on the context to achieve different cell fates. For instance, Pnt and Lozenge (Lz) binding to the *prospero* (*pros*) enhancer is required to maintain *pros* expression in R1/6/7/cone equivalence group from which R7 differentiates(18). Similarly, for cone cell expression of *shaven* (*sv*); Pnt, Lz and Suppressor of Hairless (Su(H)) binding is necessary as mutation of binding sites for any of these three factors eliminates *sv* expression in cones(22). Despite the vital roles of Pnt during development, only a few direct Pnt targets have been reported to date, including genes that direct cell type differentiation such as *pros* and *sv*. Pnt also regulates the movement of the MF by activating the expression of the morphogen *hedgehog* (*hh*). Binding of Pnt to a *hh* enhancer is required for proper *hh* expression in the eye and mutating Ets binding sites abolishes *hh* expression(21).

Chromatin Immunoprecipitation followed by next generation sequencing (ChIP-seq) is a commonly used assay to identify genome-wide occupancy of a TF. However, gene expression levels are often under the control of multiple TFs and the binding profile of a single TF is rarely sufficient to infer functional effects on transcriptional regulation. In addition, false positives frequently arise in ChIP-seq due to non-specific binding of the antibody and introduction of biases during library preparation and sequencing(23,24). Therefore, integrating ChIP-seq data with other genomic data will aid in overcoming these limitations to identify biologically relevant TF targets. Moreover, intersecting ChIP-seq with single cell genomics data sets generated from the same tissue, cell type-specific targets of TFs can be identified, which is not feasible with data derived from bulk tissues alone.

To improve our understanding of how Pnt regulates eye development and to identify its direct targets, we employed a unique approach where we integrated genome-wide data derived from different genomic assays. We first performed ChIP-seq on late third instar larval eye-antennal discs. Eye-antennal discs were dissected from animals carrying a genomic BAC DNA encoding a GFP- and FLAG-tagged Pnt protein (*pnt-GFP-FPTB; pnt*^*d88*^*/pnt*^*2*^) that fully rescues *pnt* null mutants(25). We used anti-GFP or anti-FLAG antibodies to conduct ChIP-seq and found that 6,362 and 6,268 regions (peaks) were enriched, respectively. Our ChIP-seq data show previously identified Pnt direct targets as well as many other genes whose function in the eye is currently not known. Next, we intersected our ChIP-seq data with previously published late larval eye single cell RNA-seq and ATAC-seq datasets(26) to identify 157 genes that are putative cell type-specific targets of Pnt. In addition, enhancer-reporter analyses show that our ChIP-seq data predicts functional cell type-specific enhancers that were previously unknown and may be regulated by Pnt. Together, our data is an important resource that expands the number of putative Pnt targets in the developing eye and therefore provides a platform for future studies of Pnt in development.

## Materials and Methods

### Fly stocks

The GFP- and FLAG-tagged Pnt fly stock (*w1118; PBac(pnt-GFP*.*FPTB)VK00037*) was generated using the ModEncode pipeline and is available from the Bloomington Stock Center (stock number 42680). Larval eye-antennal discs were dissected from *w*^*1118*^; *pnt-GFP-FPTB; pnt*^*Δ88*^ */ pnt*^*2*^ animals for ChIP-seq, while eye discs for immunohistochemistry were obtained from *D. melanogaster Canton-S* animals.

### Immunohistochemistry

Immunohistochemistry on larval eye discs was performed as described previously(27). Briefly, larval eye-antennal discs were dissected in 1x PBS and fixed immediately with 1xPBS+16% paraformaldehyde solution for 30 min. After washing with PBT (1xPBS+0.03% Triton-X), eye-antennal complexes were blocked in PBT supplemented with 5% normal goat serum. Primary antibody incubation was done at 4°C overnight. Secondary antibody incubation was done at room temperature for 1 to 2 hrs. The primary antibodies used are chicken anti-GFP (1:1000) and guinea pig anti-Runt (1:1000). Alexa fluorophores anti-guinea pig 647 and anti-chicken 568 were used as secondary antibodies at 1:1000 concentration.

### Chromatin immunoprecipitation

120 late third instar larval heads, containing 240 eye discs, were dissected into ice cold PBS. Dissected heads were then fixed in 1.5% formaldehyde in PBS for 15 minutes at room temperature. Fixed heads were quenched with 0.125 M glycine + PBS solution on ice for 5 minutes. Heads were washed in ice cold wash buffer A (10 mM Hepes pH 7.6, 10 mM EDTA pH 8.0, 0.5 mM EGTA pH 8.0, 0.25% Triton X-100) for 10 minutes followed by another wash in wash buffer B (10 mM Hepes pH 7.6, 100 mM NaCl, 1 mM EDTA pH 8.0, 0.5 mM EGTA pH 8.0, 0.01% Triton X-100) for 10 minutes at 4°C. Eye discs were dissected away from the heads and placed in ice cold wash buffer B. The eye discs were pelleted by centrifugation at 800 g for 30 seconds at 4°C. Eye discs were resuspended in 1 mL of sonication buffer (50 mM Hepes pH 7.6, 140 mM NaCl, 1 mM EDTA pH 8.0, 1% Triton X-100, 0.1% sodium deoxycholate, 0.1% SDS, supplemented with proteinase inhibitors (GenDepot P3100-001)) and then transferred to a 15 ml Falcon tube. Eye discs were then sonicated on ice with a Misonix S-4000 Sonicator, 3.2 mm probe. The sonication cycle was as follows: 10 s at 65 amplitude, 30 s rest on ice, total sonication time: 3 minutes. 10 µL of 10% SDS, 100 µL 1% sodium deoxycholate, 100 µL 10% Triton X-100, and 28 µL of 5M NaCl were added to the sonicated eye discs and incubated at 4°C for 10 minutes. Sonicated eye discs were then centrifuged at maximum speed for 5 minutes at 4°C to remove any debris. Supernatant containing the sonicated chromatin was transferred to a new tube. For ChIP-seq experiments, 30 µL of antibody conjugated Protein G Dynabeads were added to the supernatant and immunoprecipitated overnight at 4°C in a tube rotator. 5 µL rabbit anti-GFP (Invitrogen A-6455) and 5 µL of mouse anti-FLAG (M2, Sigma Aldrich, F1804) antibodies were used for each ChIP-seq experiment. Beads were washed once in each of the following buffers: sonication buffer, ChIP wash buffer A (same recipe as sonication buffer but with 500 mM NaCl), ChIP wash buffer B (20 mM Tris pH 8.0, 1 mM EDTA pH8.0, 250 mM LiCl, 0.5% NP-40, 0.5% sodium deoxycholate), and finally TE buffer for 5 minutes each at 4°C. To remove crosslinks, all supernatant was replaced with 150 µL of elution buffer (50 mM Tris pH 8.0, 50 mM NaCl, 2 mM EDTA, 0.75% SDS, 20 μg/mL RNase A) and incubated overnight at 65°C. Eluted chromatin was removed from the beads and set aside and 150 µL of fresh elution buffer was added to the beads and incubated at 65°C for 30 minutes. Supernatant was pooled with previously eluted chromatin to yield ∼300 µL of eluted chromatin. 60 µg of Proteinase K was added to the eluted chromatin and incubated at 37°C for 2 hours to complete the crosslink removal process. ChIP DNA was subjected to phenol-chloroform extraction and ethanol precipitation. Total input controls were treated with the same protocol as the ChIP samples, except that there were no immunoprecipitation steps. Sonicated chromatin from total input controls was directly subjected to the crosslink removal protocol followed by Proteinase K treatment, phenol chloroform extraction, and ethanol precipitation. Libraries for next generation sequencing were generated from these DNA samples.

### ChIP-seq analyses

ChIP-seq libraries were prepared and paired-end sequencing (PE75) was performed with an Illumina NextSeq 500 system. Samples were sequenced to a depth of 40 million reads. All samples passed the initial quality controls on the fastQ files. We next removed overrepresented sequences such as sequencing adapters from fastQ files using ‘cutadapt’ and ‘trimmomatic’ tools. We next aligned the reads to *Drosophila melanogaster* genome dm6 with ‘Bowtie2,’ using sensitive local alignment presets to generate a set of BAM files for the ChIP-seq data. Since we received data from four sequencing lanes (technical repeats), we merged BAM files to generate a single merged BAM file for each sample using ‘SAMtools.’ Merged BAM data were then filtered based on the MAPQ quality score such that only mapped reads that have a MAPQ quality score of at least 20 were retained. Any PCR or optical duplicates were also removed at this step. The resulting filtered BAM files were then sorted with SAMtools. MACS2 was run on the filtered and sorted BAM files using the anti-GFP or anti-FLAG ChIP as the ChIP-seq treatment file, and input control as the control file. MACS2 was run with the following parameters: effective genome size of 1.2 × 10^8^, 5 < mfold < 50, minimum qvalue cutoff of 0.01. Gene annotation of called peaks was completed using the Bedtools intersect function against all *Drosophila* genes in the dm6 reference. DNA sequences of 500 bp in length and centered on each Pnt ChIP summit were used for MEME ChIP analyses. MEME ChIP analyses were run using default parameters except for the following: expected motif distribution of zero or one occurrence per sequence, “nmotifs” of 10.

### GO analysis methods

Genes that were near the ChIP-seq peaks were used as input for analyses with Panther database.

### Cloning

Peak-region DNA was amplified by PCR and cloned into a pH-stinger-dGFP vector(28) that encodes a destabilized GFP protein. The gel extracted and cleaned peak-region DNA and vector were digested and ligated using the restriction enzymes, NheI and KpnI. Site specific integration was used to introduce the enhancer-reporter cassette into the *attP2* landing site and transgenic flies were generated by *Genetivision Corporation*. Larval eye-brain complexes from transgenic flies were used for immunohistochemistry.

## Results

### ChIP-seq binding profiles of Pnt fused to GFP and FLAG tags

To identify genes that may be regulated by Pnt, we performed ChIP-seq using late larval eye-antennal discs derived from *pnt-GFP-FPTB; pnt*^*Δ88*^*/pnt*^*2*^ animals. The *pnt-GFP-FPTB* is a transgenic genomic clone encoding a Pnt protein with GFP and FLAG tags on its C-terminus. The clone covers ∼11 kb upstream and ∼24 kb downstream of the *pnt* locus and appears to include all regulatory sequences required for proper *pnt* expression. Specifically, animals that are trans-heterozygous for *pnt* null alleles are fully rescued by the *pnt-GFP-FPTB* transgene, consistent with previous reports(29). Late larval eye-antennal discs were chosen for ChIP-seq because retinal progenitor cells are actively differentiating into different cell types at this stage of eye development. Since Pnt is required for the differentiation and survival of both neuronal and non-neuronal cells, performing ChIP-seq at this stage will allow the identification of targets that underlie these complex processes. We performed ChIP-seq with anti-GFP or anti-FLAG antibodies with two or three biological replicates, respectively. Each sample was sequenced to a depth of 40 million reads. The unique reads were then mapped to the *Drosophila melanogaster* genome release 6 (dm6), resulting in an average of 6,362 (8,787 and 3937) and 6,268 (6902, 4903 and 6998) regions (peaks) enriched by anti-GFP or anti-FLAG, respectively.

Next, peaks were mapped to genes that are within 2 kb of the peaks. An average of 7,487 genes were associated with anti-GFP ChIP-seq peaks, while anti-FLAG peaks mapped to an average of 6,396 genes. Most peaks are close to the transcription start site (average of 41.5%) while just 11.9% (anti-GFP) or 12.3% (anti-FLAG) of peaks map to intergenic regions. We also performed pairwise intersections of each possible combination of anti-GFP and anti-FLAG ChIP-seq peak sets and found overlaps ranging from 73% to 97%. Taken together, these data suggest that our ChIP-seq results are reproducible and the initial quality control is good.

### Pnt ChIP-seq identifies peaks near previously identified targets

Very few directs targets of Pnt have been identified in *Drosophila* tissues thus far, including *pros*(18), *hh*(21), *tailless* (*tll*)(30), *string* (*stg*)(19), *sv*(22) and *ETS-domain lacking* (*edl*)(31). Among these, only *pros, hh, stg* and *sv* have been identified as direct Pnt targets in the eye. Previously published analyses of enhancers in the vicinity of these genes has suggested that Pnt directly regulates their expression during development. Our ChIP-seq genomic tracks show peaks that overlap with the enhancers of these genes known to be bound and regulated by Pnt. For instance, *hh* is expressed posterior to the MF(21) (Figure 1B) and the progression of the MF requires *hh* signaling. A 203 bp minimal eye-specific enhancer in the first intron of *hh* was identified that drives reporter expression in all cells posterior to the MF, except R8. This minimal enhancer region contains Ets binding sites where Pnt binds to activate *hh* expression. All five biological repeats show a prominent Pnt-ChIP-seq peak as well as Ets binding sites that are centered on this *hh* enhancer (black box, Figure 1C). Our snATAC-seq data also show a peak in the same genomic location that aligns with the ChIP-seq peak (Figure 1D). Similarly, our data show Pnt-GFP-FPTB occupancy at a published Pnt-dependent 5’ enhancer of *pros*(18) (Supplementary Figure 1A-C) that controls expression in the R7 equivalence group, the *spa* minimal enhancer (SME) of *sv*(32) (Supplementary Figure 2A-C) that drives reporter expression in a cone-specific pattern, and the promoter of *stg* (data not shown) to activate its expression and triggering mitosis in the SMW(19). DNA gel shift assays were used to identify Ets binding sites within these enhancers bound by Pnt(18,19,32). Our snATAC-seq data also show peaks that overlap with the ChIP-seq peaks. Furthermore, Ets binding sites are also present in these enhancers suggesting that our ChIP-seq data can accurately identify previously reported Pnt binding regions and may predict other functionally relevant targets.

**Figure 1.**
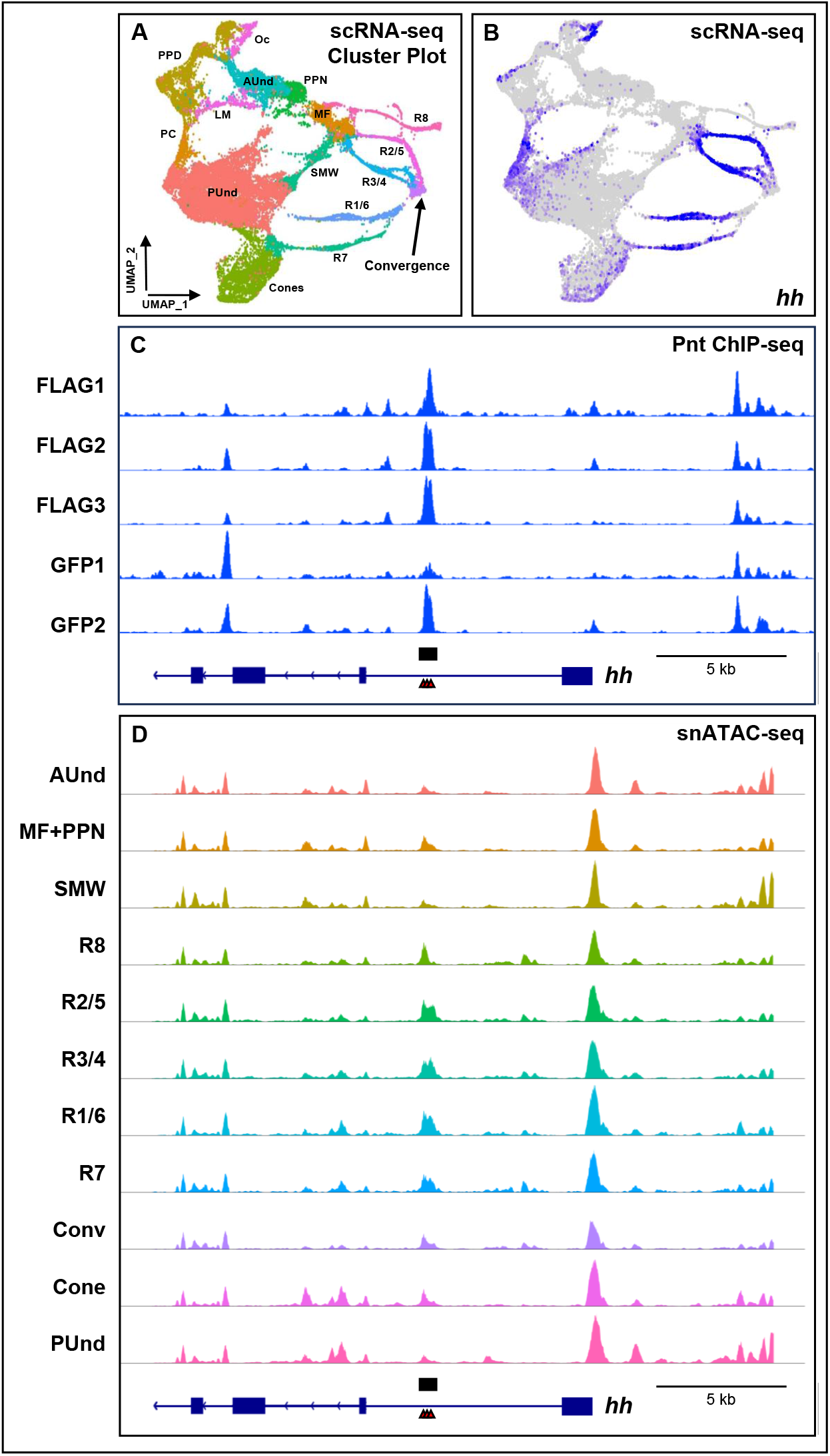
Binding profiles of Pnt ChIP-seq replicates and snATAC-seq show clear enrichment at known targets of Pnt. (A) Late larval scRNA-seq plot showing clusters corresponding to all major cell types present in the physical eye disc. (B) Plot showing the expression pattern of *hh* from single cell RNA sequencing (scRNA-seq) data. The intensity of blue is proportional to log-normalized expression levels. (C) The *hedgehog* (*hh*) locus with ChIP-seq peaks overlapping an enhancer reported to be bound by Pnt (black rectangular box). FLAG1, FLAG2, FLAG3, GFP1 and GFP2 are Pnt ChIP-seq biological replicates. (D) snATAC-seq genomic track showing the *hh* locus with peaks that overlap the known enhancer and peak shown in (C). Predicted Ets binding sites are shown as red triangles.

### Pnt binding regions are enriched for transcription factor motifs

Ets transcription factors can recruit other factors to regulate target-gene expression. For instance, the Pnt P2 isoform and Sine oculis (So) cooperatively activate *hh* and *pros* expression during eye development. We therefore used MEME-ChIP and individually subjected anti-FLAG and anti-GFP ChIP-seq peaks to motif analyses to identify binding sites of putative cofactors that may be jointly recruited along with Pnt or motifs of other transcription factors which may compete with Pnt to regulate target gene expression during eye development. Since false positives are often identified in *de novo* DNA motif analyses, we employed several approaches to filter irrelevant motifs in our analyses. First, we individually subjected each anti-GFP and anti-FLAG peak to motif analysis and identified factors that are common to at least four of the five repeats. Next, we used our scRNA-seq data to visualize the expression of the putative factors and retained only those that are expressed in the eye disc. Finally, we discarded factors that do not show overlapping expression with Pnt expression in our scRNA-seq data.

Among the top 20 motifs enriched in individual anti-FLAG and anti-GFP ChIP-seq peaks, ten motifs are common to at least four out of the five replicates (Supplementary Data 1). These are the Ets motif (Pnt and Aop), the zinc finger TF Crooked legs (Crol), the transcriptional repressors Tramtrack (Ttk) and Adult enhancer factor 1 (Aef1), the zinc finger TF Klumpfuss (Klu), the Boundary element-associated factor of 32 kD (BEAF-32), the Smad family factor Medea (Med), the Bone morphogenetic protein (BMP) signaling pathway member Mothers against dpp (Mad), the Pipsqueak type TF encoded by *CG15812*, and the paired-rule TF Paired (Prd). With the sole exception of Prd, eight of these factors are expressed in our late larval eye disc scRNA-seq dataset and show overlapping expression patterns with Pnt (Figure 2B-K). These factors are candidate binding partners of Pnt in the eye disc as they appear in all of our analyses. The remaining top 20 motifs enriched in individual ChIP-seq peaks are shown in Supplementary Data 1. As expected, the Ets motif (bound by Pnt and Anterior open/Yan) is among the top three motifs identified in anti-GFP and anti-FLAG ChIP-seq peaks of all replicates (Supplementary Data 1). It is well documented that Aop competes with Pnt for DNA binding sites to repress target gene expression(25,29). Furthermore, other transcription factors identified in our analyses have known roles in eye development and function and therefore may coregulate some target genes with Pnt(33-38).

**Figure 2.**
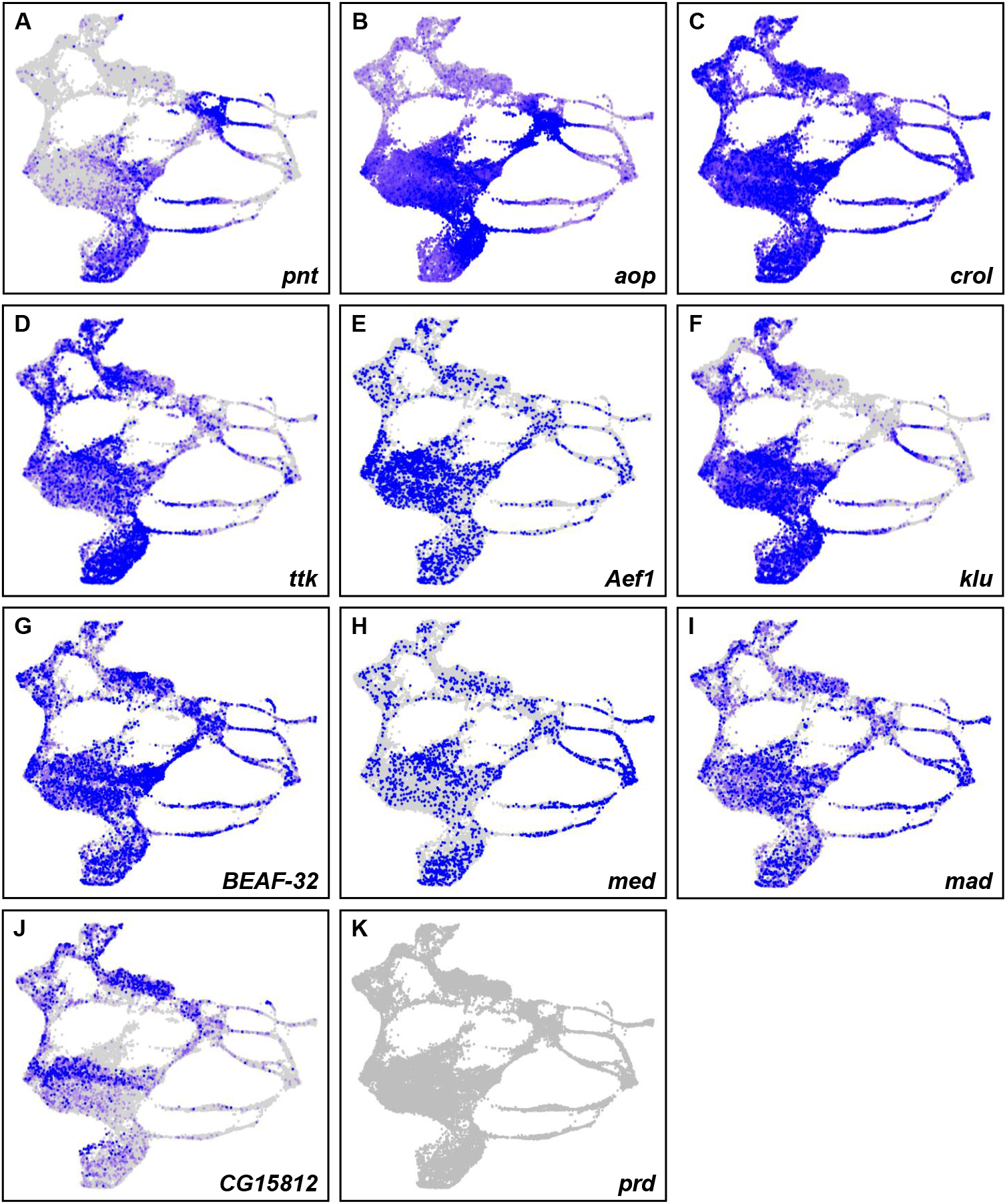
Putative coregulators of Pnt are expressed in overlapping patterns with *pnt*. (A-K) scRNA-seq plots showing the expression patterns of *pnt* (A), *aop* (B), *crol* (C), *ttk* (D), *Aef1* (E), *klu* (F), *BEAF-32* (G), *med* (H), *mad* (I), *CG15812* (J), and *prd* (K). Other than *prd*, all genes show overlapping expression with *pnt*.

### Pnt ChIP-seq peaks near genes involved in eye function

To identify the biological processes that may be regulated by Pnt, we performed Gene Ontology (GO) analyses with genes associated with the ChIP-seq peaks using the Panther database(39). As expected, some of the top enriched GO terms are associated with processes related to eye function and development (Supplementary aDta 2). These include the *sevenless* (*sev*) signaling pathway (GO:0045501), regulation of cell cycle G1/S phase transition (GO:1902806), compound eye cone cell differentiation (GO:0042675), R3/R4 cell fate commitment (GO:0007464), positive regulation of cell cycle G1/S phase transition (GO:1902808), imaginal disc growth (GO:0007446), negative regulation of photoreceptor cell differentiation (GO:0046533) and positive regulation of dendrite morphogenesis (GO:0050775). GO analyses also identified several processes that are related to EGFR and MAP kinase signaling. These include ERBB signaling pathway (GO:0038127), regulation of phosphatidylinositol 3-kinase signaling (GO:0014066) and epidermal growth factor receptor signaling pathway (GO:0007173) Processes that are related to other signaling pathways including the cytokine-mediated signaling pathway (GO:0019221) and the hippo mediated signaling pathway (GO:0035332) were identified by GO analyses. Finally, we also see GO clusters that are unrelated to eye function, such as eggshell chorion gene amplification (GO:0007307), wing and notum subfield formation (GO:0035309), and male anatomical structure morphogenesis (GO:0090598). This may be expected because Pnt is known to be expressed and activate several targets in a wide range of *Drosophila* tissues(10,11,14,17). Taken together, these data identify many genes that may be regulated by Pnt during eye development that were previously unknown targets.

### Identification of putative cell type-specific targets of Pnt

The Egfr pathway is responsible for the sequential differentiation of both neuronal photoreceptors (except R8) and non-neuronal cone and pigment cells. The mechanism by which reiterative use of Egfr triggers these different outcomes within the eye disc is not well understood. One hypothesis is that depending on the cell state (i.e., the transcriptional milieu and/or chromatin accessibility), distinct targets may be activated in different cell types upon Egfr pathway induction. To identify putative cell type-specific targets of Pnt, we intersected our ChIP-seq data with single nuclear ATAC-seq (snATAC-seq) and single cell RNA sequence (scRNA-seq) datasets generated from late larval eye discs(26),(25,29). We employed several criteria to identify novel putative direct targets of Pnt in the eye: 1) the gene has not been previously reported as a cell type-specific Pnt target; 2) a ChIP-seq peak maps within 2 kb of the gene; 3) a snATAC-seq peak overlaps the ChIP-seq peak that maps to the gene; and 4) the gene shows cell type-specific expression in the scRNA-seq dataset. We intersected single cell genomics datasets with anti-GFP and anti-FLAG ChIP-seq datasets; this yielded 157 or 145 genes in the anti-GFP or anti-FLAG ChIP-seq datasets, respectively, that are specifically expressed in R1-7 or cone cells (Supplementary Data 3-5). All 145 anti-FLAG genes are present in anti-GFP gene list. Motif analyses using peaks that map to these 145 genes from anti-GFP and anti-FLAG repeats identified six factors that appear in at least four out of the five replicates: Ets, CroI, Aef1, Klu, Lame duck (Lmd) and Buttonhead (Btd) (Supplementary Data 6). Although our scRNA-seq data suggests that Lmd and Btd are not expressed at detectable levels in the eye (not shown), the other four factors are expressed in the eye and overlap with *pnt* expression (Figure 2). Supplementary Data 6 shows a complete list of factors (with E-values) identified in the individual repeat analyses.

Among these 145 unique genes, 127 of the genes that map to anti-GFP peaks and 135 of the genes associated with anti-FLAG peaks show multiple Ets binding sites (two or more) within 200 bp of the peak summit. These genes include many putative novel targets of Pnt during eye development and several are known to be involved in retinal cell type specification. For example, the *rough* (*ro*) gene is expressed in the MF, R2/5 and R3/4(40) (Figure 3B) and is required for the proper specification of R2 and R5. The first intron harbors a *ro* enhancer that drives reporter expression in the MF, R2/5 and R3/4 (black box, Figure 3A). All five ChIP-seq repeats show a peak in the first intron of *ro* that overlaps with a snATAC-seq peak at the same genomic location (Figure 3C). The peak region also contains four Ets binding sites, suggesting that *ro* is a cell type-specific target of Pnt. Similarly, *spalt major* (*salm*), *sevenless* (*sev*), and *seven up* (*svp*) (Supplementary Figures 3-5) are putative novel Pnt targets that are involved in cell type specification in the eye and also show overlapping Pnt ChIP-seq and snATAC-seq peaks. While some genes in these lists have no reported eye function, a few genes with known roles in axon development and function are present. In addition, 47 anti-GFP and 46 anti-FLAG genes are expressed in a cell type-specific manner in late larval eye discs (Supplementary Data 5). Moreover, 42 genes are common to both datasets and a total of 51 unique genes are identified when both GFP and FLAG gene lists are combined. One example is the Fibroblast growth factor (FGF) pathway member *pyramus* (*pyr*), which is predominantly expressed in R1/6 and R7 (Figure 4B). All five ChIP-seq repeats show a prominent peak ∼10 kb downstream of *pyr* (Figure 4A). The snATAC-seq genomic track also shows a peak in the same genomic location that is most accessible in R1/6 and R7 and a cluster of Ets binding sites are present in this peak region (Figure 4C). This suggests that *pyr* may be a cell type-specific target of Pnt. To test if this peak contains a cell type-specific functional enhancer, we analyzed the peak region DNA fragment *in vivo* for enhancer activity. We amplified the peak-region DNA of *pyr* (black box, Figure 4B,C), cloned it in front of a destabilized green florescent protein (dGFP) encoding reporter gene, and generated transgenic flies carrying this construct. Larval eye discs of these flies were stained with GFP and Runt (an R7 and R8 marker) antibodies (Figure 4D-F). We observe that GFP colocalizes with Runt in some but not all ommatidial clusters. The GFP and Runt expressing cell is apical to a second cell that expresses only Runt. This arrangement reflects the known positions of R7 and R8 cells in the developing eye, suggesting that GFP expression is in the R7 cell. Taken together, these results suggest that our ChIP-seq data can predict functional and cell type-specific enhancers in the eye. Other putative novel cell type-specific targets of Pnt, including *factor of interpulse interval* (*fipi*) (Supplementary Figure 6A-C), are shown in Supplementary Data 5.

**Figure 3.**
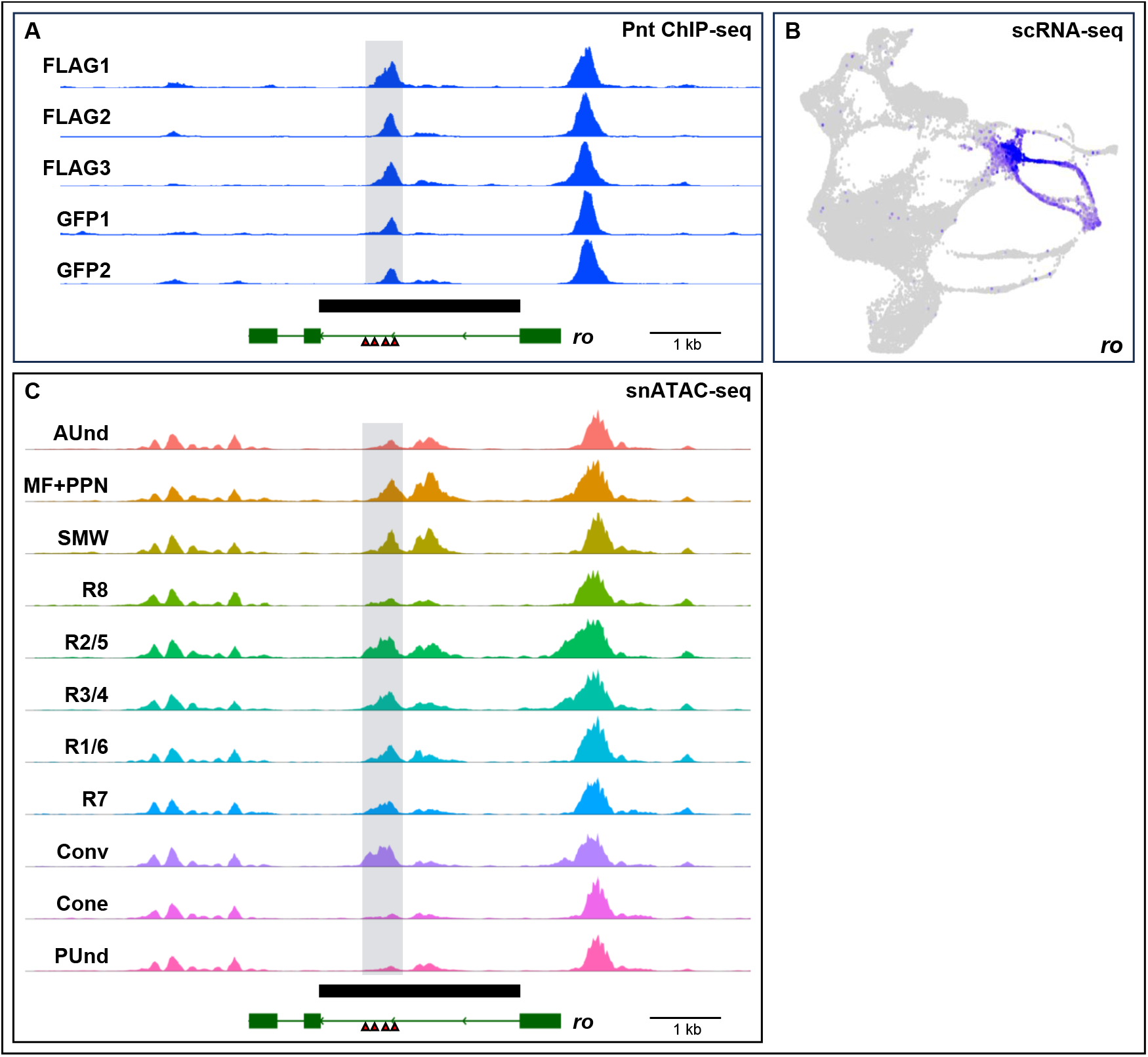
Pnt ChIP-seq peak profiles are found near genes with known essential roles in eye development. (A) ChIP-seq genomic tracks showing peaks in the intron of *rough* (*ro*). The peak region is highlighted in gray. The solid black bar indicates a known enhancer of *ro*. (B) scRNA-seq plot showing the expression of *ro* in the MF, R2/5 and R3/4. (C) snATAC-seq plot showing the *ro* locus with a peak in an intron that aligns with the ChIP-seq peak. The peak region is highlighted in gray. Four Pnt binding sites are shown as red triangles.

**Figure 4.**
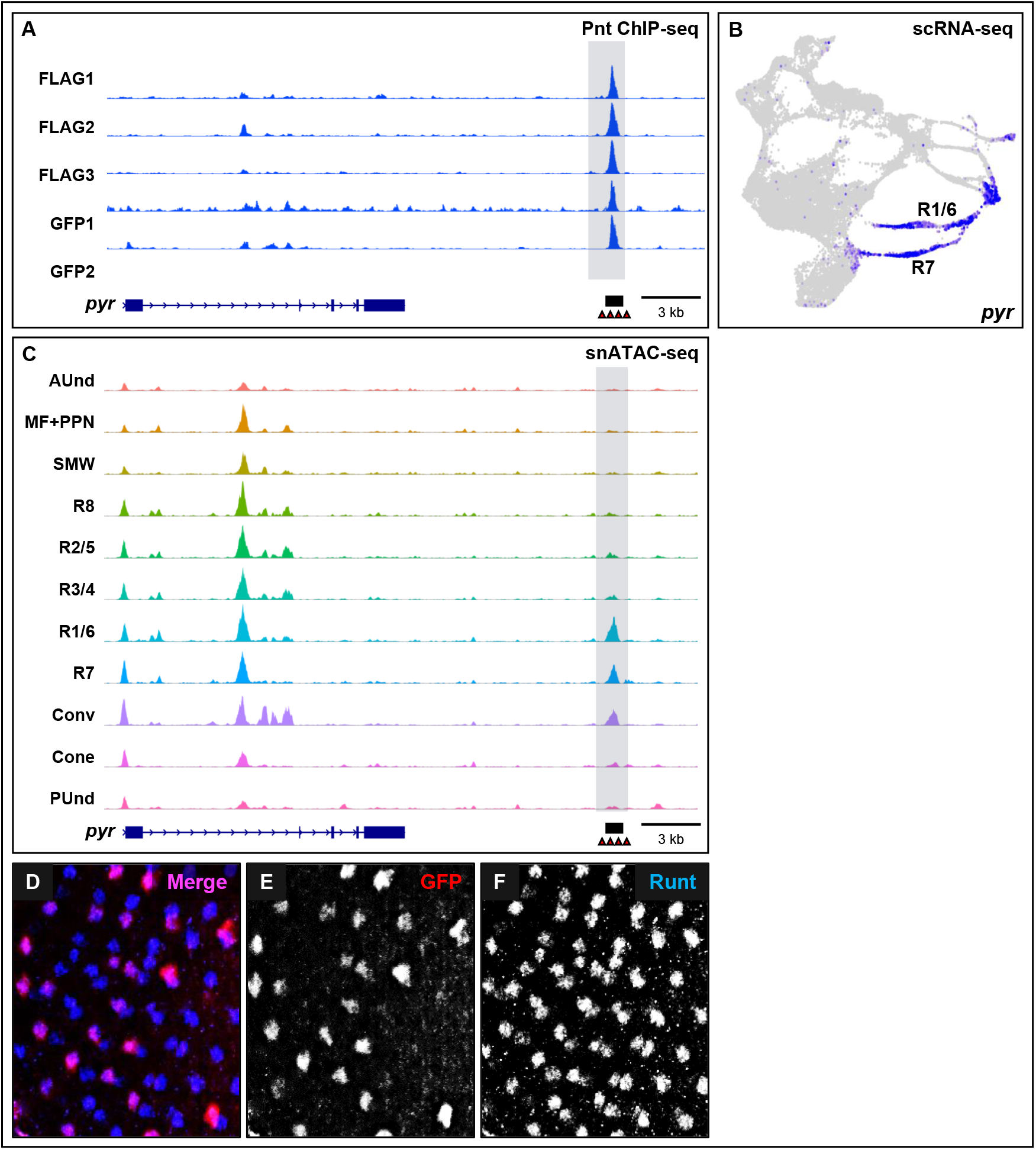
*pyramus* is a putative novel cell type-specific target of Pnt. (A) Genomic track showing a prominent Pnt ChIP-seq peak about 11 kb 3’ of *pyramus* (*pyr*). (B) Larval eye disc scRNA-seq plot showing the expression (blue) of *pyr* in R1/6, R7 and the Convergence cell clusters. (C) snATAC-seq genomic tracks at the *pyr* locus show prominent peaks in R1/6, R7 and the Convergence cell clusters. This peak is present at the same genomic location as the 3’ peak in the ChIP-seq data sets (A). The black box represents the peak-region DNA that was tested for enhancer activity *in vivo*. (D-F) Immunostaining of transgenic larval eye discs carrying the peak-region DNA in (A) and (C) driving expression of a dGFP reporter. Larval eye discs were stained with GFP (E) and Runt (F), which marks R7 and R8. Many ommatidia show coexpression of GFP and Runt antibodies (D).

## Discussion

The Ets domain transcription factor Pnt is vital for *Drosophila* eye development, and yet very few direct Pnt targets are currently known. We used ChIP-seq to identify the genome-wide binding profile of Pnt in late larval eye discs. Our data show an average of 6,362 and 6,268 binding regions enriched by anti-GFP or -FLAG antibodies, respectively, with an overlap of about 84%. These regions map to an average of 7,487 and 6,396 genes, respectively, and include previously known Pnt targets as well as many novel putative targets that may be relevant to eye development. Intersecting our ChIP-seq with previously published single cell datasets reveals putative cell type-specific enhancers and targets regulated by Pnt. We present an expanded list of putative Pnt targets that may play important roles in larval eye disc development.

The Ets family of transcription factors are functionally versatile and regulate the expression of a variety of genes in diverse tissues and cell types. This functional adaptability of Ets factors may reflect their interactions with other transcription factors that enable combinatorial control of gene expression. Motif analyses using the ChIP-seq peaks identified in this study reveals putative Pnt-interacting transcription regulators in the eye. As predicted, the Ets motif, to which either Pnt or Aop can bind, is one of the top motifs found in Pnt ChIP-seq peaks. The role of Aop as a transcriptional repressor is well documented. Aop competes with Pnt for Ets binding sites and opposes the gene activation by Pnt. In some tissues, Pnt and Aop show mutually exclusive expression patterns with varying levels of Pnt and Aop driving differentiation of cell types by competing for the same binding sites(25,29). Similarly, antagonistic action of Pnt and the transcriptional repressor Ttk is required for the correct transcription of the cell cycle gene *stg*, which controls proliferation of cells in the SMW(19). Fine tuning of gene expression by Ets factors and their antagonistic repressors may be a common mechanism to confer robustness of target gene expression during development.

Our ChIP-seq peak datasets also show enrichment for the putative zinc finger transcription factor Crol, which is activated by ecdysone-regulated gene expression. This suggests a possible link between Pnt and the hormone ecdysone, which triggers larval molting, pupariation, as well as MF progression and the cell cycle in the eye disc(35,41). Furthermore, *crol* regulates cell cycle progression in *Drosophila* wing discs. Since Pnt plays vital roles in cell cycle and MF progression, it is possible that Crol is a cofactor of Pnt in regulating these events in the eye. Future functional studies that modulate the activities of Pnt and Crol should unravel their significance in cell cycle progression and regulation.

Mad is another factor identified in our motif screen to identify Pnt coactivators. Mad acts downstream of the *decapentaplegic* (*dpp*) pathway and is involved in the initiation of the MF and may have a minor role in MF propagation(34). Furthermore, Mad is known to interact with the cofactor Med(36). Since Pnt activates *hh* and plays a role in MF propagation, it is possible that Pnt, Mad and Med may cooperatively regulate this event in the eye. Future studies will be needed to more fully understand the functional significance of these and several other motifs identified in our Pnt ChIP-seq dataset.

How Egfr activation of Pnt leads to the differentiation of several different cell types is currently unknown. Pnt may cooperate with other factors and regulate expression of downstream targets by a combinatorial mechanism. Under this model, unique cell fates are achieved depending on the binding partner of Pnt. For example, *pros* and *sv* are direct targets of Pnt in the eye. The R7/cone equivalence group comprises the precursor cells of R7 and the cones and *pros* expression is limited to these precursors by Pnt and the transcription factor Lz. Similarly, expression of *sv* in cones requires Pnt, Lz, and Su(H), which is a downstream effector of the Notch signaling pathway. Binding site mutations of any one of these factors abolishes cone cell expression driven by the *sv* enhancer. Therefore, activation of Pnt, context-dependent cofactors, and a unique cell type response collectively appear to underlie the differentiation of different cell types in the eye disc. By overlapping our ChIP-seq data with late larval single cell genomics datasets, we identified target genes that show cell type-specific expression patterns. These genes may be mediators of the unique cell type responses resulting from activation of Pnt and may play critical roles in retinal cell differentiation. Analyzing loss-of-function mutants will unravel the function of these genes in cell fate decisions in the eye.

In summary, we report a high quality ChIP-seq dataset that expands the list of putative Pnt direct targets that may be involved in the differentiation of individual cell types in the eye. Our data therefore represents an important resource for researchers studying *Drosophila* eye development as well as studies examining the diverse roles of Ets transcription factors in regulating their downstream targets.

## Data availability

All raw data are uploaded onto Gene Expression Omnibus, Accession number: XXXXXX.

## Acknowledgements

We thank Claude Desplan for sharing anti-Runt antibodies. Library prep and sequencing was performed at the Genomic and RNA Profiling Core at Baylor College of Medicine.

## Author Contributions

K.K.B.R. and K.Y. performed the larval eye disc dissection. The ChIP-seq protocol and analyses were performed by K.Y. K.K.B.R. prepared and conducted the immunofluorescence imaging of larval eye discs. Y.S. performed enhancer cloning. The manuscript was prepared by K.K.B.R. and reviewed by K.Y. and G.M.

## Correspondence

Correspondence should be sent to Graeme Mardon.

## References

1. Wasylyk, B., Hahn, S.L. and Giovane, A. (1993) The Ets family of transcription factors. European journal of biochemistry, 211, 7–18.

2. Oikawa, T., Yamada, T., Kihara-Negishi, F., Yamamoto, H., Kondoh, N., Hitomi, Y. and Hashimoto, Y. (1999) The role of Ets family transcription factor PU. 1 in hematopoietic cell differentiation, proliferation and apoptosis. Cell Death & Differentiation, 6, 599–608.

3. Maroulakou, I.G. and Bowe, D.B. (2000) Expression and function of Ets transcription factors in mammalian development: a regulatory network. Oncogene, 19, 6432–6442.

4. Hollenhorst, P.C., McIntosh, L.P. and Graves, B.J. (2011) Genomic and biochemical insights into the specificity of ETS transcription factors. Annual review of biochemistry, 80, 437–471.

5. Dittmer, J. and Nordheim, A. (1998) Ets transcription factors and human disease. Biochimica et Biophysica Acta-Reviews on Cancer, 1377, F1.

6. Oikawa, T. (2004) ETS transcription factors: possible targets for cancer therapy. Cancer science, 95, 626–633.

7. Seth, A. and Watson, D.K. (2005) ETS transcription factors and their emerging roles in human cancer. European journal of cancer, 41, 2462–2478.

8. Turner, D.P. and Watson, D.K. (2008) ETS transcription factors: oncogenes and tumor suppressor genes as therapeutic targets for prostate cancer. Expert review of anticancer therapy, 8, 33–42.

9. Hsing, M., Wang, Y., Rennie, P.S., Cox, M.E. and Cherkasov, A. (2020) ETS transcription factors as emerging drug targets in cancer. Medicinal research reviews, 40, 413–430.

10. Scholz, H., Deatrick, J., Klaes, A. and Klämbt, C. (1993) Genetic dissection of pointed, a Drosophila gene encoding two ETS-related proteins. Genetics, 135, 455–468.

11. Hsu, T. and Schulz, R.A. (2000) Sequence and functional properties of Ets genes in the model organism Drosophila. Oncogene, 19, 6409–6416.

12. Mayer, U. and Nüsslein-Volhard, C. (1988) A group of genes required for pattern formation in the ventral ectoderm of the Drosophila embryo. Genes & development, 2, 1496–1511.

13. Jan, Y.N. and Jan, L.Y. (1994) Genetic control of cell fate specification in Drosophila peripheral nervous system. Annual review of genetics, 28, 373–393.

14. Klaes, A., Menne, T., Stollewerk, A., Scholz, H. and Klämbt, C. (1994) The Ets transcription factors encoded by the Drosophila gene pointed direct glial cell differentiation in the embryonic CNS. Cell, 78, 149–160.

15. Schottenfeld, J., Song, Y. and Ghabrial, A.S. (2010) Tube continued: morphogenesis of the Drosophila tracheal system. Current opinion in cell biology, 22, 633–639.

16. O’Neill, E.M., Rebay, I., Tjian, R. and Rubin, G.M. (1994) The activities of two Ets-related transcription factors required for Drosophila eye development are modulated by the Ras/MAPK pathway. Cell, 78, 137–147.

17. Freeman, M. (1996) Reiterative use of the EGF receptor triggers differentiation of all cell types in the Drosophila eye. Cell, 87, 651–660.

18. Xu, C., Kauffmann, R.C., Zhang, J., Kladny, S. and Carthew, R.W. (2000) Overlapping activators and repressors delimit transcriptional response to receptor tyrosine kinase signals in the Drosophila eye. Cell, 103, 87–97.

19. Baonza, A., Murawsky, C.M., Travers, A.A. and Freeman, M. (2002) Pointed and Tramtrack69 establish an EGFR-dependent transcriptional switch to regulate mitosis. Nature cell biology, 4, 976–980.

20. Yang, L. and Baker, N.E. (2003) Cell cycle withdrawal, progression, and cell survival regulation by EGFR and its effectors in the differentiating Drosophila eye. Developmental cell, 4, 359–369.

21. Rogers, E.M., Brennan, C.A., Mortimer, N.T., Cook, S., Morris, A.R. and Moses, K. (2005) Pointed regulates an eye-specific transcriptional enhancer in the Drosophila hedgehog gene, which is required for the movement of the morphogenetic furrow. Development, 132, 4833–4843.

22. Rohrbaugh, M., Ramos, E., Nguyen, D., Price, M., Wen, Y. and Lai, Z.-C. (2002) Notch activation of yan expression is antagonized by RTK/pointed signaling in the Drosophila eye. Current biology, 12, 576–581.

23. Park, P.J. (2009) ChIP–seq: advantages and challenges of a maturing technology. Nature reviews genetics, 10, 669–680.

24. Liu, P., Sanalkumar, R., Bresnick, E.H., Keleş, S. and Dewey, C.N. (2016) Integrative analysis with ChIP-seq advances the limits of transcript quantification from RNA-seq. Genome research, 26, 1124–1133.

25. Webber, J.L., Zhang, J., Massey, A., Sanchez-Luege, N. and Rebay, I. (2018) Collaborative repressive action of the antagonistic ETS transcription factors Pointed and Yan fine-tunes gene expression to confer robustness in Drosophila. Development, 145.

26. Raja, K.K.B., Yeung, K., Shim, Y.-K., Li, Y., Chen, R. and Mardon, G. (2023) A Single Cell Genomics Atlas of the <em>Drosophila</em> Larval Eye Reveals Distinct Developmental Timelines and Novel Markers for All Photoreceptor Subtypes. bioRxiv, 2023.2002.2014.528565.

27. Hsiao, H.-Y., Johnston Jr, R.J., Jukam, D., Vasiliauskas, D., Desplan, C. and Rister, J. (2012) Dissection and immunohistochemistry of larval, pupal and adult Drosophila retinas. JoVE (Journal of Visualized Experiments), e4347.

28. Pepple, K.L., Atkins, M., Venken, K., Wellnitz, K., Harding, M., Frankfort, B. and Mardon, G. (2008) Two-step selection of a single R8 photoreceptor: a bistable loop between senseless and rough locks in R8 fate.

29. Boisclair Lachance, J.F., Peláez, N., Cassidy, J.J., Webber, J.L., Rebay, I. and Carthew, R.W. (2014) A comparative study of Pointed and Yan expression reveals new complexity to the transcriptional networks downstream of receptor tyrosine kinase signaling. Dev Biol, 385, 263–278.

30. Chen, R., Deng, X. and Zhu, S. (2022) The Ets protein Pointed P1 represses Asense expression in type II neuroblasts by activating Tailless. PLoS Genet, 18, e1009928.

31. Vivekanand, P., Tootle, T.L. and Rebay, I. (2004) MAE, a dual regulator of the EGFR signaling pathway, is a target of the Ets transcription factors PNT and YAN. Mechanisms of Development, 121, 1469–1479.

32. Flores, G.V., Duan, H., Yan, H., Nagaraj, R., Fu, W., Zou, Y., Noll, M. and Banerjee, U. (2000) Combinatorial signaling in the specification of unique cell fates. Cell, 103, 75–85.

33. Xiong, W.C. and Montell, C. (1993) tramtrack is a transcriptional repressor required for cell fate determination in the Drosophila eye. Genes & development, 7, 1085–1096.

34. Wiersdorff, V., Lecuit, T., Cohen, S.M. and Mlodzik, M. (1996) Mad acts downstream of Dpp receptors, revealing a differential requirement for dpp signaling in initiation and propagation of morphogenesis in the Drosophila eye. Development, 122, 2153–2162.

35. D′ Avino, P.P. and Thummel, C.S. (1998) crooked legs encodes a family of zinc finger proteins required for leg morphogenesis and ecdysone-regulated gene expression during Drosophila metamorphosis. Development, 125, 1733–1745.

36. Das, P.L. Maduzia, L., Wang, H. L. Finelli, A., Cho, S.-H., Smith, M.M. and Padgett, R.W. (1998) The Drosophila gene Medea demonstrates the requirement for different classes of Smads in dpp signaling. Development, 125, 1519–1528.

37. Rusconi, J.C., Fink, J.L. and Cagan, R. (2004) klumpfuss regulates cell death in the Drosophila retina. Mechanisms of development, 121, 537–546.

38. Jukam, D., Viets, K., Anderson, C., Zhou, C., DeFord, P., Yan, J., Cao, J. and Johnston, R.J., Jr. (2016) The insulator protein BEAF-32 is required for Hippo pathway activity in the terminal differentiation of neuronal subtypes. Development, 143, 2389–2397.

39. Mi, H., Lazareva-Ulitsky, B., Loo, R., Kejariwal, A., Vandergriff, J., Rabkin, S., Guo, N., Muruganujan, A., Doremieux, O. and Campbell, M.J. (2005) The PANTHER database of protein families, subfamilies, functions and pathways. Nucleic acids research, 33, D284–D288.

40. Tomlinson, A., Kimmel, B.E. and Rubin, G.M. (1988) rough, a Drosophila homeobox gene required in photoreceptors R2 and R5 for inductive interactions in the developing eye. Cell, 55, 771–784.

41. Abrell, S., Carrera, P. and Jäckle, H. (2000) A modifier screen of ectopic Krüppel activity identifies autosomal Drosophila chromosomal sites and genes required for normal eye development. Chromosoma, 109, 334–342.

