## Supplementary_Figures for "Integrative Genomic Analyses Reveal Putative Cell Type-specific Targets of the *Drosophila* Ets Transcription Factor Pointed"

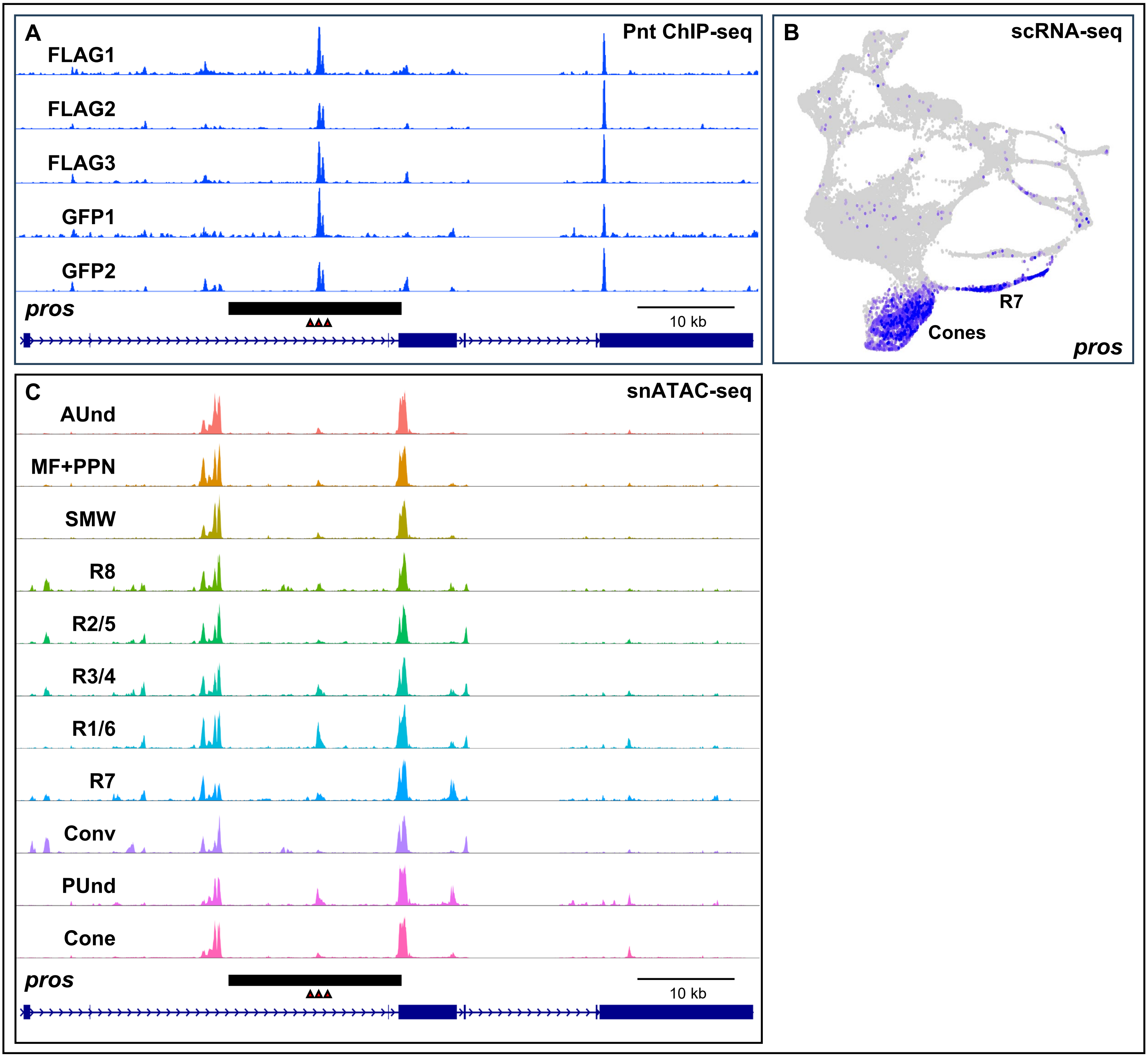

A
Pnt ChIP-seq
FLAG1
FLAG2
FLAG3
GFP1
GFP2
pros
10 kb
B
scRNA-seq
R7
Cones
pros
C
snATAC-seq
AUnd
MF+PPN
SMW
R8
R2/5
R3/4
R1/6
R7
Conv
PUnd
Cone
pros
10 kb

### Supplementary Figure Legends

**Supplementary Figure 1.** Binding profiles of Pnt ChIP-seq replicates show clear peaks within the *prospero* locus, a known target of Pnt. (A) ChIP-seq genomic tracks of *prospero* (*pros*) an enhancer known to be regulated by Pnt. The enhancer is shown as a black rectangular box. FLAG1, FLAG2, FLAG3, GFP1 and GFP2 are the ChIP-seq biological repeats. (B) scRNA-seq plot showing the expression pattern of *pros*. (C) snATAC-seq genomic track showing *pros* gene locus with peaks that overlap the known enhancer. Ets binding sites are shown as red triangles in all Supplementary Figures.

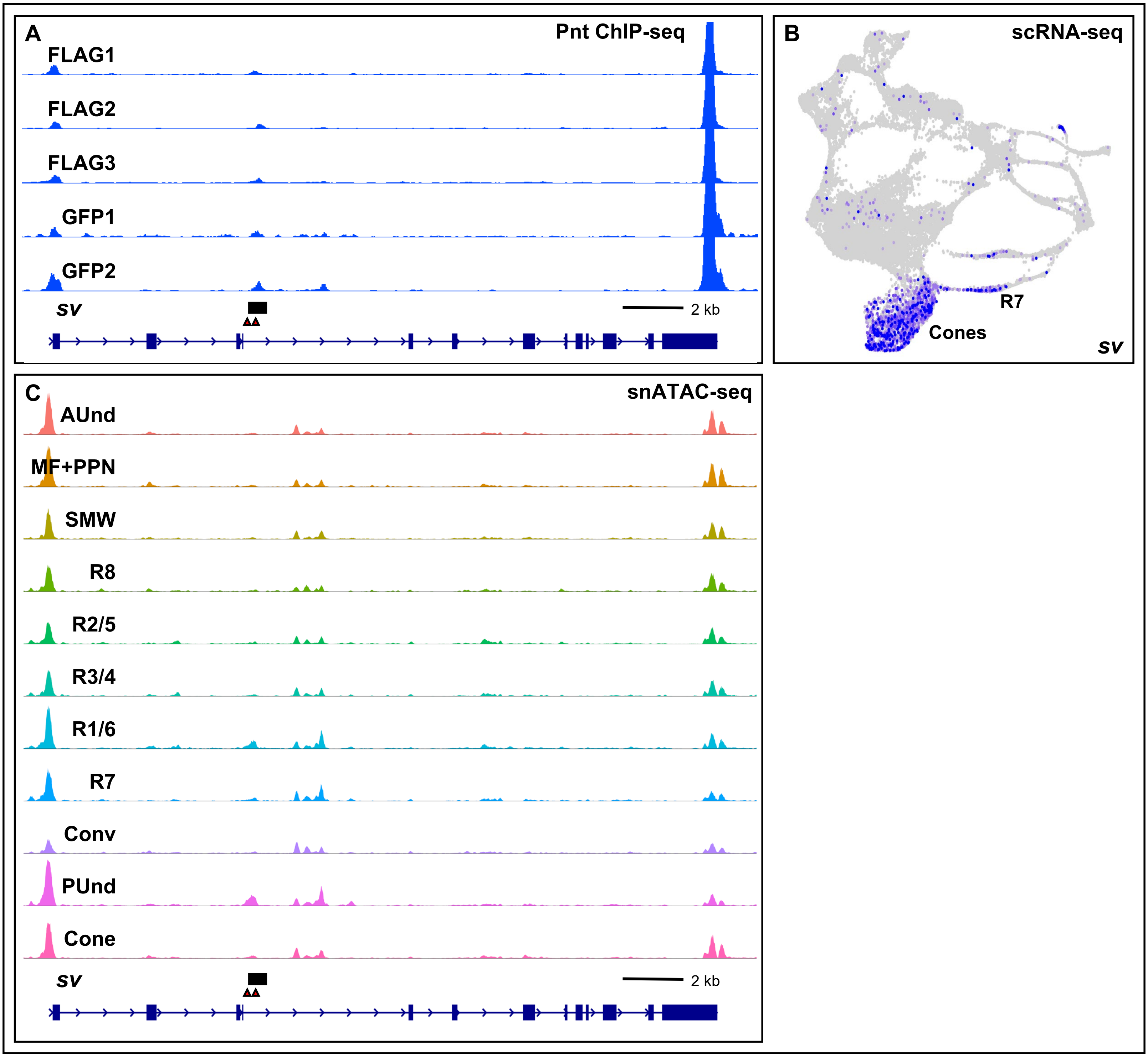
A
Pnt ChIP-seq
FLAG1
FLAG2
FLAG3
GFP1
GFP2
sv
2 kb
B
scRNA-seq
R7
Cones
sv
C
snATAC-seq
AUnd
MF+PPN
SMW
R8
R2/5
R3/4
R1/6
R7
Conv
PUnd
Cone
sv
2 kb

**Supplementary Figure 2.** Binding profiles of Pnt ChIP-seq replicates show clear peaks near the *shaven* locus, a known target of Pnt. (A) ChIP-seq genomic tracks of *shaven* (*sv*) showing an enhancer regulated by Pnt. The enhancer is shown as a black rectangular box. (B) scRNA-seq plot showing the expression pattern of *sv* in R7 and cone cells. (C) snATAC-seq track showing the *sv* locus with peaks that overlap the known enhancer.

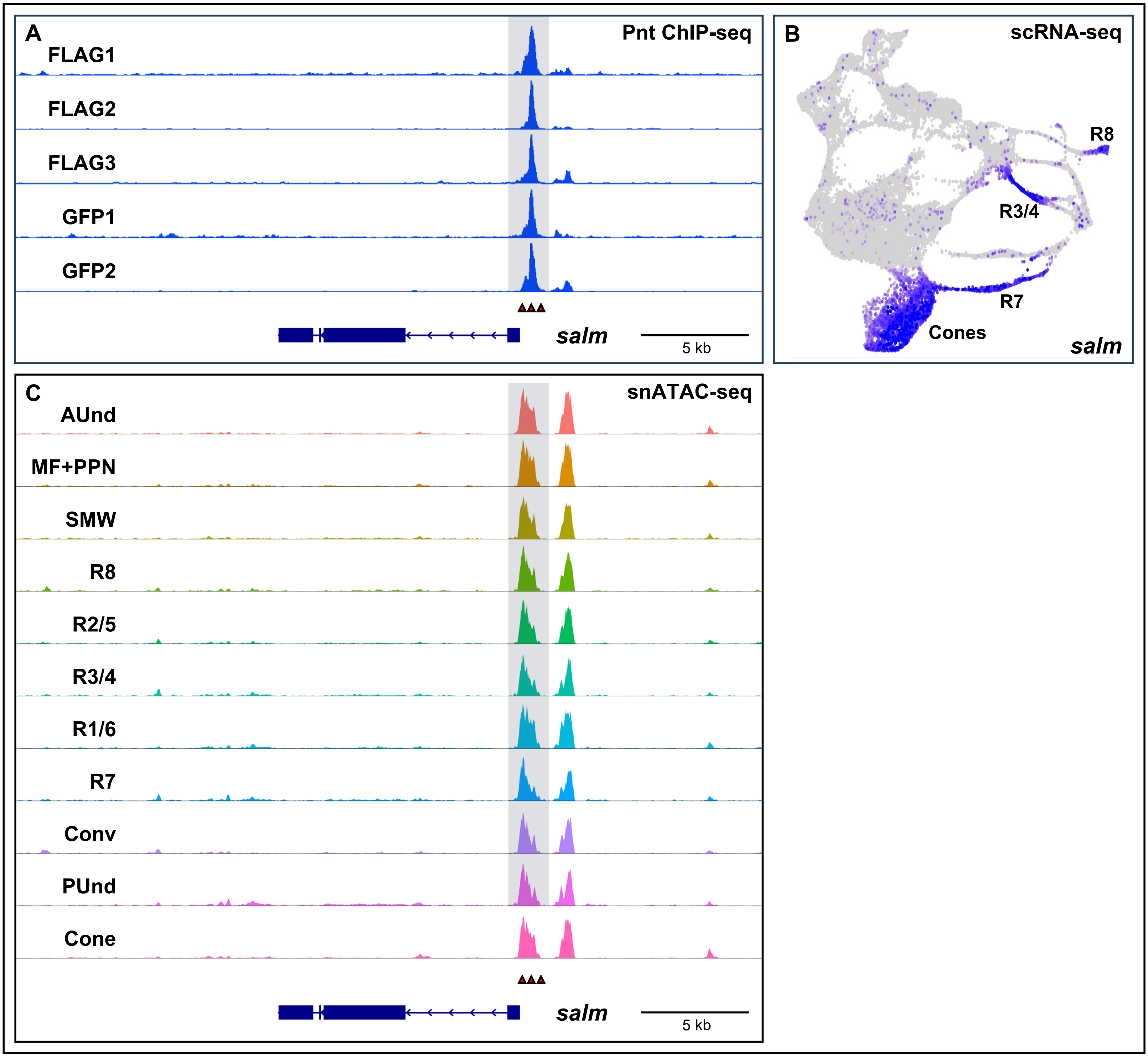

A
Pnt ChIP-seq
FLAG1
FLAG2
FLAG3
GFP1
GFP2
salm
5 kb
B
scRNA-seq
R8
R3/4
R7
Cones
salm
C
snATAC-seq
AUnd
MF+PPN
SMW
R8
R2/5
R3/4
R1/6
R7
Conv
PUnd
Cone
salm
5 kb

**Supplementary Figure 3.** Pnt ChIP-seq peaks near *spalt major*, a known regulator of eye development. (A) ChIP-seq genomic tracks of *spalt major* (*salm*) showing a peak immediately upstream of the transcription start site. (B) Late larval scRNA-seq plot showing *salm* expression in R3/4, R7, R8 and cone cells. (C) snATAC-seq genomic track showing a peak that overlaps the ChIP-seq peaks.

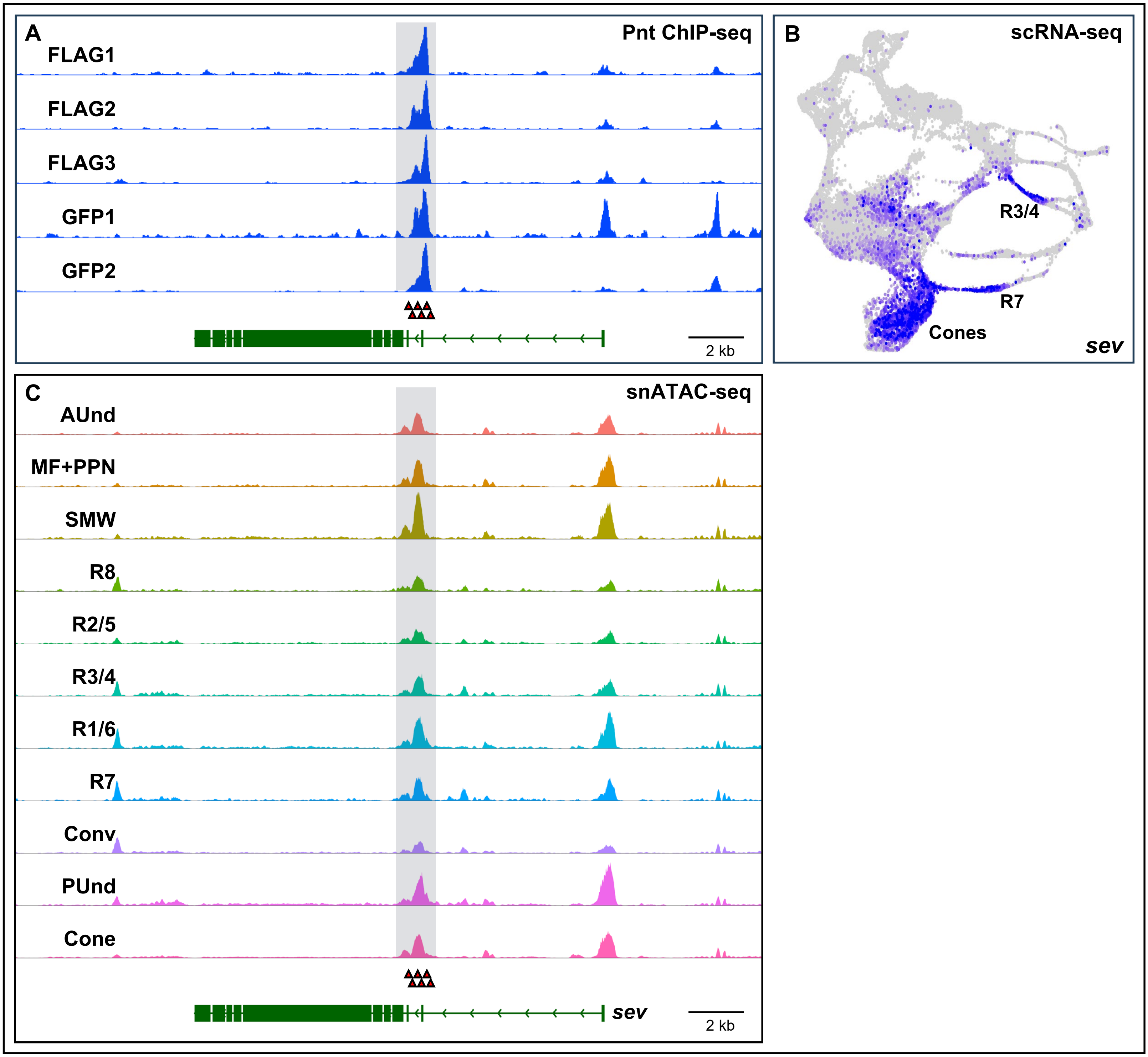
A
Pnt ChIP-seq
FLAG1
FLAG2
FLAG3
GFP1
GFP2
2 kb
B
scRNA-seq
R3/4
R7
Cones
sev
C
snATAC-seq
AUnd
MF+PPN
SMW
R8
R2/5
R3/4
R1/6
R7
Conv
PUnd
Cone
sev
2 kb

**Supplementary Figure 4.** Pnt ChIP-seq peaks within the *sevenless* locus, a known regulator of R7 development. (A) ChIP-seq genomic tracks of *sevenless* (*sev*) showing a peak overlapping the second exon. (B) Late larval scRNA-seq plot showing *sev* expression. *sev* is predominantly expressed in R3/4, R7 and cone cells. (C) snATAC-seq genomic track showing a peak that overlaps the ChIP-seq peaks in (A).

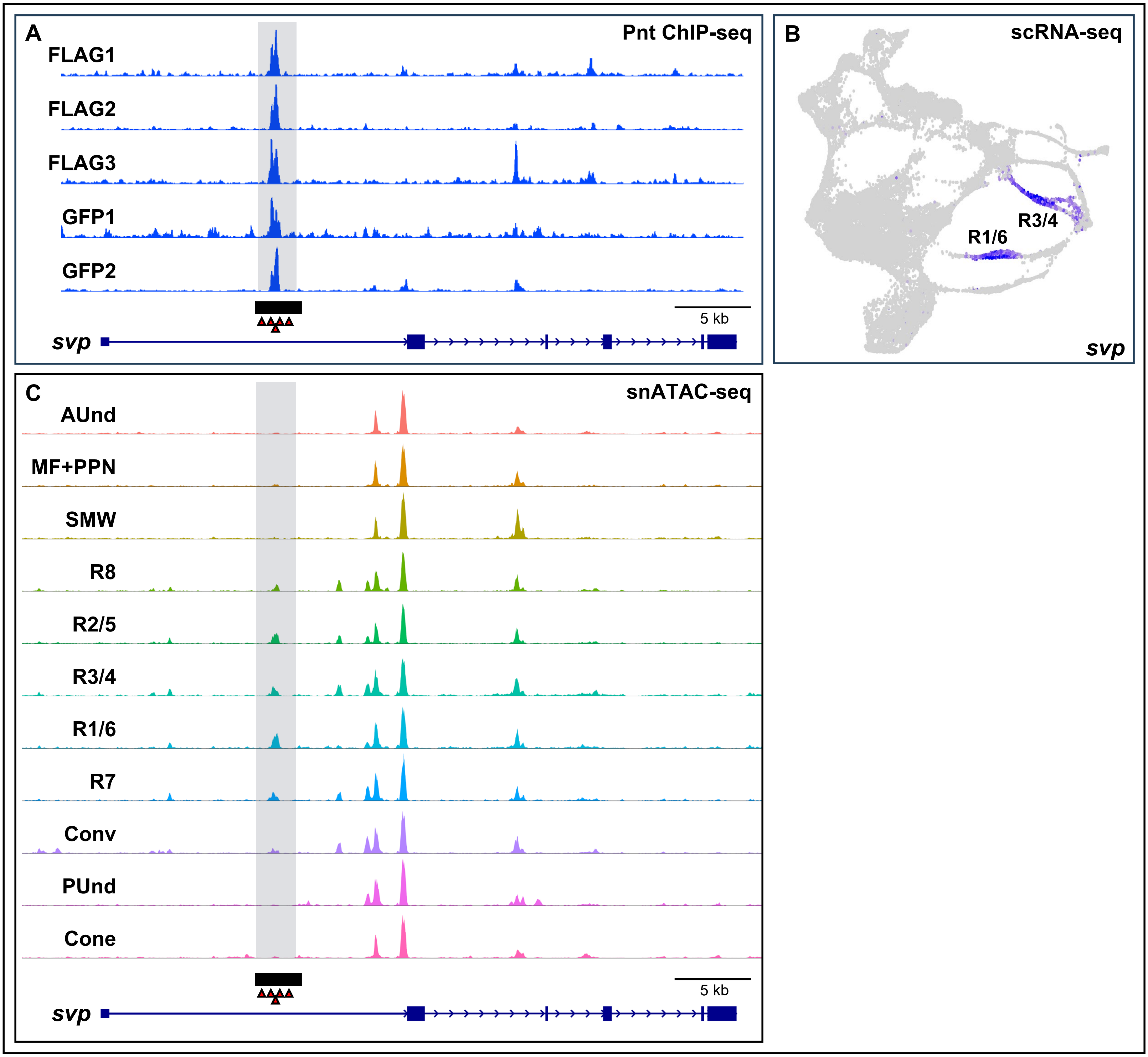
A
Pnt ChIP-seq
FLAG1
FLAG2
FLAG3
GFP1
GFP2
5 kb
svp
B
scRNA-seq
R3/4
R1/6
svp
C
snATAC-seq
AUnd
MF+PPN
SMW
R8
R2/5
R3/4
R1/6
R7
Conv
PUnd
Cone
5 kb
svp

**Supplementary Figure 5.** Pnt ChIP-seq peaks within the *seven up* locus, a known regulator of eye development. (A) ChIP-seq genomic tracks of *seven up* (*svp*) showing a peak in the first intron. The black bar represents the known *svp* enhancer. (B) Late larval scRNA-seq plot showing *svp* expression in R3/4 and R1/6. (C) snATAC-seq genomic track of the *svp* locus showing a peak that overlaps the ChIP-seq peaks.

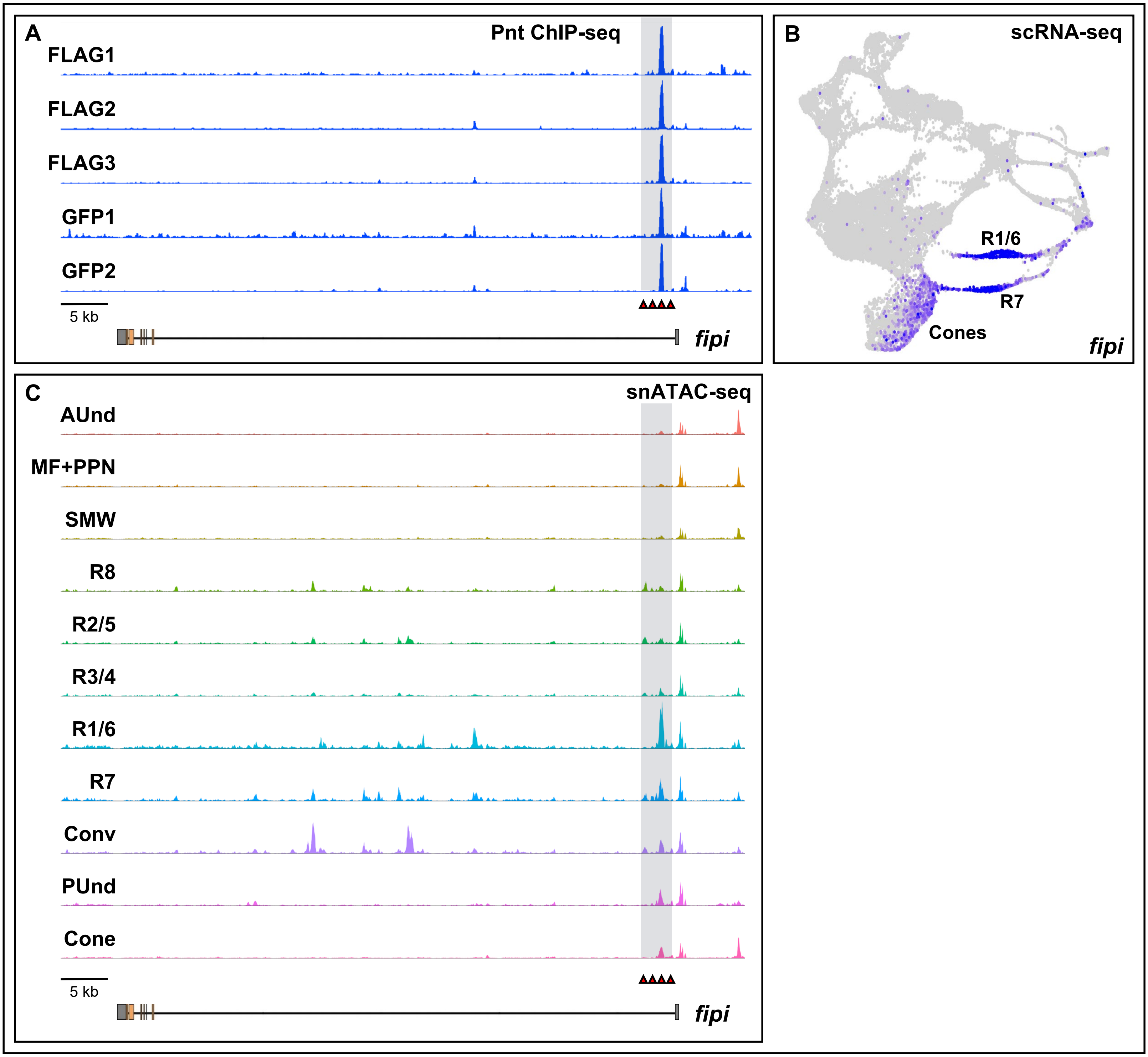
A
Pnt ChIP-seq
FLAG1
FLAG2
FLAG3
GFP1
GFP2
5 kb
fipi
B
scRNA-seq
R1/6
R7
Cones
fipi
C
snATAC-seq
AUnd
MF+PPN
SMW
R8
R2/5
R3/4
R1/6
R7
Conv
PUnd
Cone
5 kb
fipi

**Supplementary Figure 6.** *factor of interpulse interval* (*fipi*) is a putative novel cell type-specific Pnt target. (A) Pnt ChIP-seq genomic tracks show a prominent peak early in the first intron of *fipi*. (B) Larval eye disc scRNA-seq plot showing the expression pattern of *fipi* in R1/6/7 and cone cells. (C) snATAC-seq genomic tracks showing peaks that overlap with the ChIP-seq peaks shown in (A). The peak shows greater accessibility in R1/6 compared to other cell types.
